# Environmental cues shape extracellular vesicles biogenesis and function in *Streptococcus pneumoniae*

**DOI:** 10.1101/2025.07.18.665510

**Authors:** Miriana Battista, Yann Bachelot, Teresa Franke, Christoph Saffer, Lioba Zimmermann, Laura Teuber, Marc Thilo Figge, Cláudia Vilhena

## Abstract

The Gram-positive human pathogen *Streptococcus pneumoniae* adapts its metabolism to the environment during colonization and host invasion. Extracellular vesicles (EVs) are produced by *S. pneumoniae* in the process of infection but the exact interplay between metabolic adaptation and vesicle formation remains elusive. Here, we demonstrate that exposure to normal human serum induced rearrangement of the pneumococcal cell wall and significantly increased Sp-EVs production. Temperature and pH were critical factors for Sp-EVs formation: 37°C supported optimal EV production, while bacterial exposure to either basic or acidic environments slowed down pneumococcal EV biogenesis and led to a heterogeneous subpopulation profile. Proteomic analysis revealed that Sp-EVs are enriched in carbon metabolism-related proteins, specially those associated with glycolysis (e.g. eno, gapA, gapN, gpmA, pfkA, pykF, and tpi). Low glucose availability enhanced Sp-EVs production and intracellular ATP level, underlying a relation between metabolic status and EV biogenesis. Functionally, Sp-EVs promoted biofilm formation in both *S. pneumoniae* and *Streptococcus pyogenes*. Sp-EVs isolated under glucose-rich conditions enhanced *S. pneumoniae* biofilms, whereas Sp-EVs from glucose-poor conditions strongly stimulated *S. pyogenes* biofilm formation. These findings underscore the role of host and environmental cues in shaping pneumococcal EV production, composition, and function, highlighting their potential involvement in metabolic adaptation and interspecies interactions.

**Importance:** *Streptococcus pneumoniae* remains a leading cause of morbidity and mortality worldwide, responsible for diseases ranging from community-acquired pneumonia to meningitis and sepsis. It is of great concern in children under the age of five and in low-income countries, despite the existence of vaccination. The success of this pathogen relies on its capacity to adapt to diverse host niches, including the blood, nasopharynx and the lungs. Bacterial extracellular vesicles (EVs) have emerged as mediators of virulence and communication, yet their regulation by environmental and metabolic factors remains poorly understood. By exploring how environmental conditions and nutrient availability shape pneumococcal EV production and how these vesicles contribute to biofilm formation and interspecies interactions, our study contributes to the understanding of how *S. pneumoniae* applies complex strategies to persist and spread, with potential future applications in infection control and therapeutic strategies.

## Introduction

Bacteria constantly adapt to changing environmental conditions in order to survive and propagate^1,2^. A set of metabolic, genetic and general regulatory rearrangements take place, shaping the course of bacterial evolution^3–6^. For instance, bacteria can evolve to acquire antibiotic resistance^7^ or to specialize in the consumption of a specific source^8^. Environmental factors such as temperature, pH, osmolarity, and nutrient availability influence bacterial physiology^9–12^. When in contact with the human host, bacteria experience a wide array of stimuli which are sensed by surface exposed receptors and can trigger stress responses that impact bacterial fitness and hence survival^12–14^. Such stimuli range from immune molecules, antimicrobial peptides, cellular surfaces, hormones, reactive oxygen species, etc^15,16^.

*Streptococcus pneumoniae* is a Gram-positive human pathogen and the causal agent of complicated diseases as community-acquired pneumonia, meningitis and sepsis^17,18^. In *S. pneumoniae*, metabolic adaptation plays a key role in colonization and pathogenicity. For instance, during colonization of the nasopharynx, glucose and hyaluronic acid sources are prioritized^19,20^, ATP levels required for capsular polysaccharide synthesis are regulated^21^ and endogenous hydrogen peroxide (H2O2) is produced via SpxB to mitigate oxidative stress^22^. Another event that occurs during changing environmental conditions and/or host exposure is the production of extracellular vesicles (EVs)^23^.

EVs are non-replicative membrane-enclosed structures, ranging in size from 30 to 300 nm and originate from all kingdoms of life including microbial organisms ^23–26^. They contain proteins, lipids, RNA and DNA and play relevant physiological roles such as waste disposal, membrane remodeling and cell-to-cell communication^27,28^. Besides their physiological role, bacterial EVs can be implicated in, e. g. biofilm formation, cellular damage and immune modulation of the host ^29–32^. Specifically, *S. pneumoniae* EVs (Sp-EVs) have been shown to modulate host cytokine and chemokine expression and release despite their non-cytotoxic feature^33–38^. Moreover, Sp-EVs downregulate expression of an endothelial surface marker and are internalized by human endothelial cells^38^. In our previous work, we utilized a co-cultivation (bacterial-human) multi-well model to investigate whether different stimuli, e. g. host-released substances, influenced vesicle synthesis^38^. We observed a co-cultivation time and concentration-dependent EV formation which suggests that host and pneumococcal EVs biogenesis are not independent mechanisms but appear to mutually influence each other in a highly dynamic and multifaceted manner.

Despite advances in understanding the environmental and metabolic regulation of bacterial pathogenesis, the interplay between EV formation, bacterial metabolism, and their functional roles in infection remains largely unexplored. How metabolic shifts and environmental signals shape EV biogenesis, and how this, in turn, influences bacterial survival, biofilm formation, and inter-species communication, remains an open question.

Here, we further characterized the role of human serum, temperature and pH on Sp-EVs biogenesis. We found glycolysis and general carbon metabolism pathways enriched in EVs proteome. A deeper dissection of glucose influence on Sp-EVs revealed a correlation between sugar abundance and EVs concentration which relates to cellular ATP content. Finally, we tested the functional role of glucose-derived EVs and conclude that Sp-EVs formed under high sugar abundance led to distinct bacterial growth phenotypes and intra- and inter-species biofilm formation.

## Results

### Pneumococcal cell wall arrangement and Sp-EVs formation are affected by human serum

Bacteria cell wall composition and EVs production can be influenced by different environmental conditions and host-derived factors such as serum components^39,40^. Thus, we investigated the interaction of *S. pneumoniae* D39 with normal human serum (NHS) and visualized bacterial cell morphology by super-resolution microscopy (**Fig. 1A**). Mid-exponential phase D39 cells were incubated with NHS and later stained with fluorescently labeled wheat germ agglutinin (WGA), which is a carbohydrate-binding lectin with high affinity for N-acetylglucosamine (a main structural component of the pneumococcal cell wall), and DAPI for DNA.

**Fig. 1.**
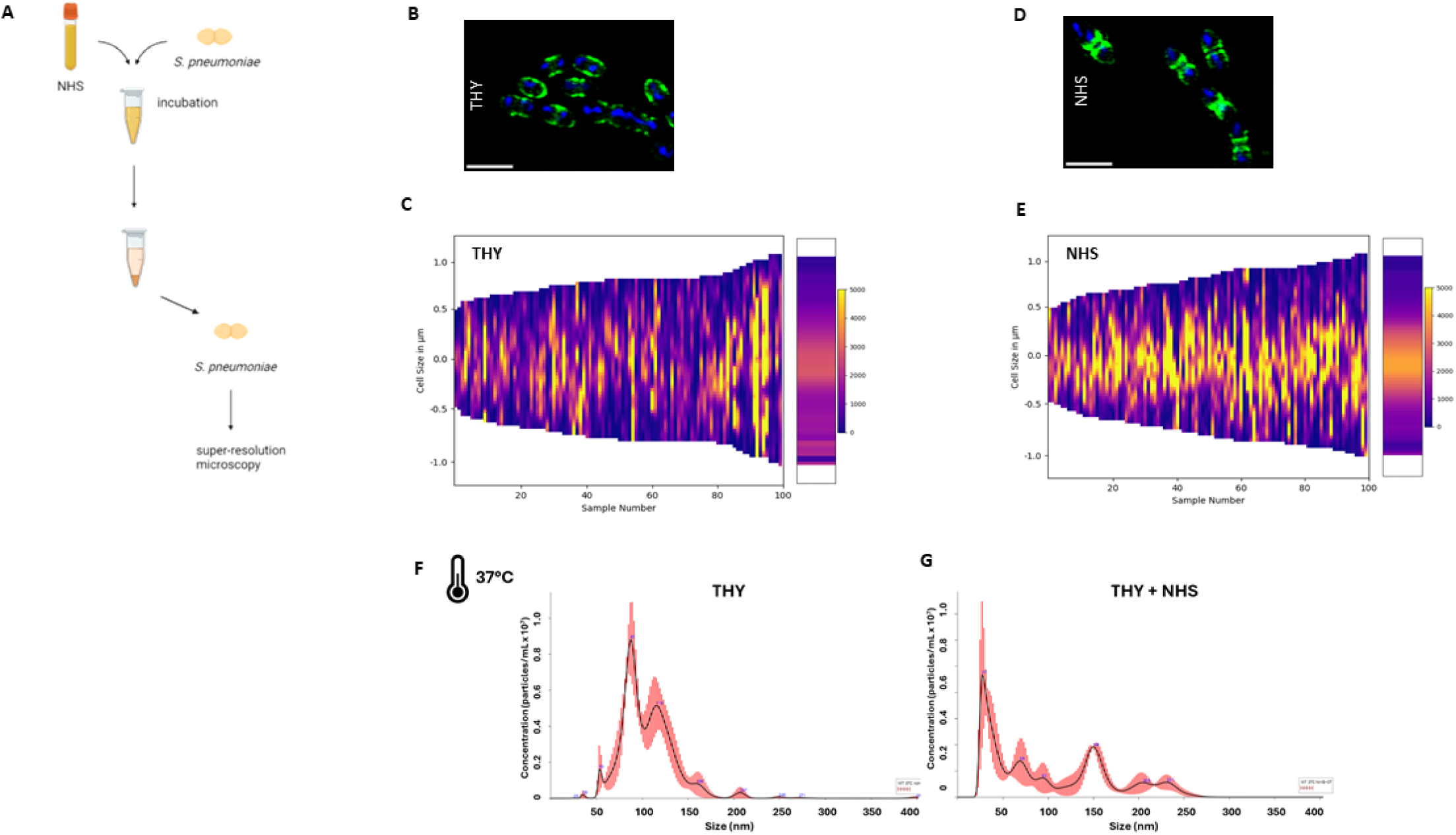
E**f**fect **of normal human serum on *S. pneumoniae* D39 cell wall and Sp-EV production.** (**A**) Schematic representation of the experiment performed to investigate the interaction of *S. pneumoniae* D39 with normal human serum (NHS). Mid-exponentially grown D39 bacterial cells were incubated with NHS for 30 minutes, centrifuged, stained and visualized by super-resolution structured illumination microscopy (SR-SIM). Representative SR-SIM images of pneumococci stained with WGA CF^®^488 (green) and DAPI (blue) in (**B**) growth medium (THY, control condition) and (**D**) after incubation with NHS. Scale bar = 2 µm (**C, E**) Graphs showing distribution of WGA across the bacterial cell wall in the control condition (THY) and in NHS-challenged bacterial cells, respectively. (**F, G**) Histograms representing Sp- EVs concentration (particles/mL × 10^7^) and size (nm), after nanoparticle tracking analysis (NTA). EVs were isolated from supernatant of *S. pneumoniae* D39 grown until mid-logarithmic phase in THY at 37°C with or without NHS.

The distribution of WGA across the bacterial cell wall in the control condition (cells growing in THY) was random and dispersed across the entire wall, as seen in the representative microscopy picture and on the distribution graph (**Fig. 1B, 1C**). A certain degree of heterogeneity among single individual bacteria could be observed, however without a correlation between cell length and signal intensity. For bacterial cells challenged with NHS, the signal distribution was different, and WGA appeared mainly localized at mid-cell, suggesting a cell wall rearrangement compared to the control (**Fig. 1D**). Quantification of signal intensity corroborated the findings, revealing a higher accumulation of WGA at the septum and flanking-septum regions in bacterial cells under NHS challenge (**Fig. 1E**).

In a previous work, we have documented the effect of human factors on pneumococcal EVs (Sp-EVs) formation, by co-cultivating human endothelial and bacterial cells^38^. Given the effect of NHS on the bacterial cell wall arrangement, we wondered if NHS would affect EV formation. To ascertain the influence of NHS on EVs formation, pneumococci were grown in the absence and presence of NHS at 37°C, EVs were isolated and further characterized by nanoparticle tracking analysis (NTA). Histograms representing particles profile showed a different size distribution and concentration of EVs isolated from NHS-challenged bacteria compared to the control (**Fig. 1F, 1G**). EV population of the control condition depicted a typical size distribution, with one main population around 80-100 nm (**Fig. 1F**). Under NHS challenge, the EVs populations appeared more heterogeneous, with several subpopulations emerging in the size range 50-150 nm (**Fig. 1G**).

Taken together, these results suggest a different cell wall arrangement of *S. pneumoniae* D39 and a modulation of EV formation when encountering human serum factors.

### Sp-EVs production is sensitive to temperature and pH variation

We showed that pneumococcal EVs production was modulated by the presence of NHS. Building on this observation, we investigated the influence of additional environmental factors such as temperature and pH.

Consistently, our results showed that Sp-EVs concentration was severely affected by distinct environmental conditions (**Fig. 2A**) but not the average size of the population (**Fig. 2B)**. Sp-EVs production was stimulated by the presence of NHS, but temperature emerged as a critical factor for this process. When pneumococci were grown at 37°C, Sp-EVs concentration in the bacterial supernatant was high (1.93 x 10^8^ particles/mL) and further increased under NHS condition (5.53 x 10^8^ particles/mL). In contrast, a temperature of 30°C led to reduced EVs concentration despite the presence of NHS (5.10 x 10^7^ particles/mL) (**Fig. 2A**). A similar size profile was observed for Sp-EVs isolated from pneumococci grown at 30°C with or without NHS (**Fig. 2C, 2D**).

**Fig. 2.**
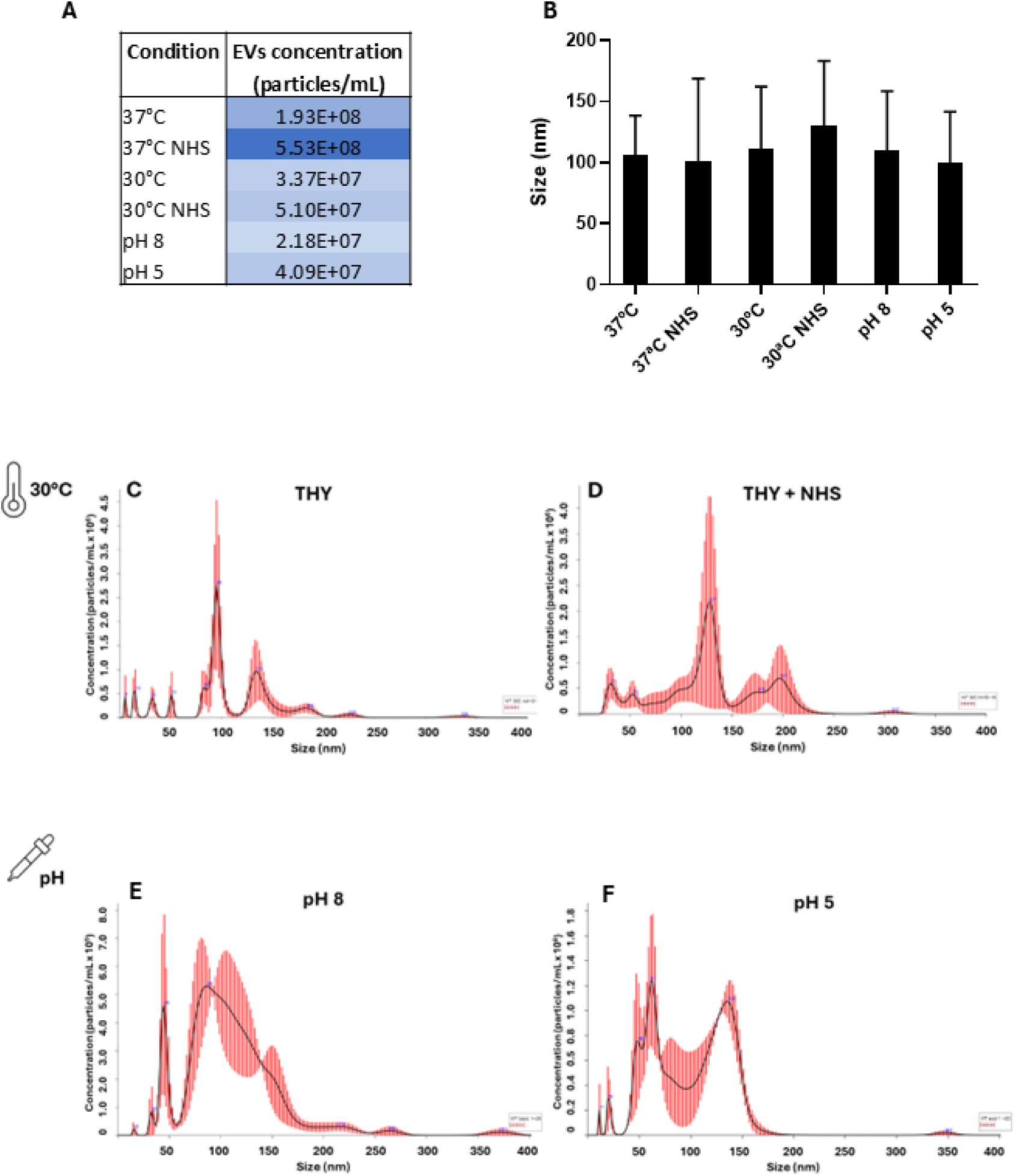
C**h**aracterization **of Sp-EVs produced under varied environmental conditions.** (**A**) Table representing Sp-EVs concentration (particles/mL) and (**B**) graph showing Sp-EVs size (nm) after nanoparticle tracking analysis. EVs were isolated from the supernatant of *S. pneumoniae* D39 grown until mid-logarithmic phase in THY at 37°C or 30°C with or without NHS or at different pH conditions. Mean ± SD. (**C-F**) Histograms showing particle profile for size (nm) and concentration (particle/mL) of EVs isolated from bacterial culture at 30°C (**C**) in the absence or (**D**) in the presence of NHS and (**E-F**) at different pH conditions (pH 8 and 5).

Additionally, variations on pH drastically affected Sp-EVs production. EVs concentration decreased when pneumococci were exposed to either basic or acidic environment (pH 8 and pH 5) in comparison with the THY control (**Fig. 2A**). The number of subpopulations increased with the alkalinity of the media (**Fig. 2E**) which goes in line with the previously observed effect of aggregation of EVs at acidic pH, leading to a wider and single subpopulation^41^.

In summary, our findings demonstrate that a temperature of 37°C is the optimal condition for *S. pneumoniae* to produce EVs. Exposure to either basic or acidic environments, as well as temperature shifts, negatively affects Sp-EVs production. In this context, environmental conditions emerge as key aspects to consider in the study of Sp-EVs biogenesis.

### Carbon metabolism-related proteins are enriched in Sp-EVs cargo

We showed that Sp-EVs production is influenced by environmental factors such as the presence of human serum, temperature and pH variations. Bacterial EVs cargo is, in turn, dynamically adapted according to environmental conditions^42^.

In our previous work, proteomic profiling on EVs isolated from *S. pneumoniae* D39 and clinical isolated strains revealed a diverse protein composition, ranging from choline-binding proteins, cell division-related proteins, capsular polysaccharide biosynthesis, etc^38^. However, this analysis primarily focused on the differential protein abundance between strains (wild-type vs clinical isolated) and did not address the functional relationships or potential co-associations between proteins. To bridge this gap and understand the functional significance and interaction networks of Sp-EVs proteins, we constructed a protein-protein interaction (PPI) map using STRING^43^ (**Fig. 3A**).

**Fig. 3.**
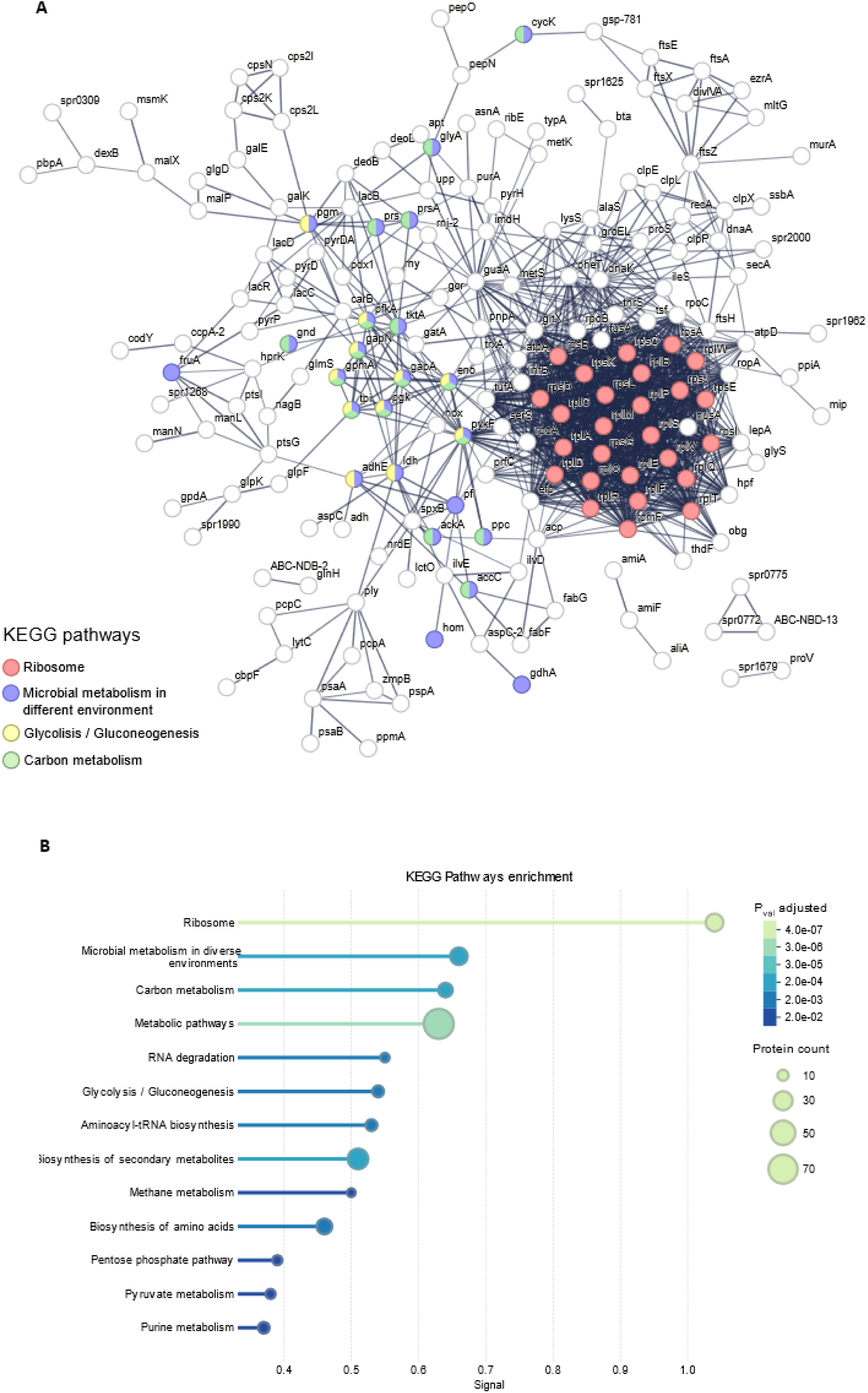
Protein-protein interaction network and enrichment analysis of Sp-EVs proteome. (**A**) STRING network of the proteins found in Sp-EVs in the proteomic analysis. Nodes and edges represent respectively the different proteins and the proteins’ interactions with high confidence (interaction score > 0.7). Proteins involved in pathways of interest are highlighted with different colors. (**B**) Functional enrichment of the network against the KEGG database. Horizontal bars show the –log10 of the adjusted p-value for every pathway that is significantly over-represented (p ≤ 0.05). Circle area is proportional to the number of Sp-EV proteins assigned to the corresponding pathway.

Functional annotation through KEGG pathway enrichment showed significantly enriched pathways in which Sp-EVs proteins are involved. Among the enriched pathways with higher signal, we found *ribosome*, *microbial metabolism in diverse environments*, *carbon metabolism*, *metabolic pathways*, *RNA degradation* and *glycolysis/ gluconeogenesis* (**Fig. 3B**).

The presence of ribosome-related proteins in Sp-EVs is in accordance with previous observations for EVs isolated from other Gram-positive and Gram-negative bacteria^44,45^. Interestingly, a substantial subset of Sp-EVs proteins mapped to core metabolic pathways, specifically to carbon metabolism and glycolysis or gluconeogenesis (e.g., eno, gapA, gapN, gpmA, pfkA, pykF, and tpiA). Eno, gapA and tpiA are known to bind human plasmin and plasminogen and to contribute to invasion ^46–48^.

These findings suggest possible involvement of Sp-EVs in mechanisms of bacterial metabolic adaptation, potentially facilitating survival and persistence under changing environmental conditions.

### Sp-EVs production is promoted by low glucose availability

PPI analyses of the proteins found in Sp-EVs cargo revealed enrichment in carbon metabolism-related proteins, several of which are involved in glycolysis or gluconeogenesis pathways. Glucose metabolism is known to regulate pneumococcal virulence properties such as capsule synthesis, rapid growth in the bloodstream, adherence and immune evasion in the nasopharynx ^49–52^. Pneumococcal extracellular vesicles, in turn, play a role in infection mechanisms and have been shown to carry virulence factors ^38,53,54^. Bringing these concepts together, we wondered if glucose availability would impact bacterial capacity to synthesize EVs.

To investigate the possible connection between glucose metabolism and Sp-EVs formation, pneumococci were grown in THY with increasing concentrations of glucose (0,10, 50, 100 and 200 mM) and Sp-EVs production was assessed.

Sp-EVs were produced in more abundance when bacteria were exposed to a lower glucose concentration (10 mM), followed by a marked decrease in total vesicle amount with increasing glucose concentrations (50, 100, 200 mM) (**Fig. 4A**). This inverse relationship suggests that EV biogenesis may be sensitive to metabolic status. Representative light scattering microscopy images (**Fig. 4B**) and particle profile (**Fig. 4C**) of Sp-EVs isolated under glucose abundance showed a homogenous morphology and a unimodal sized-population.

**Fig. 4.**
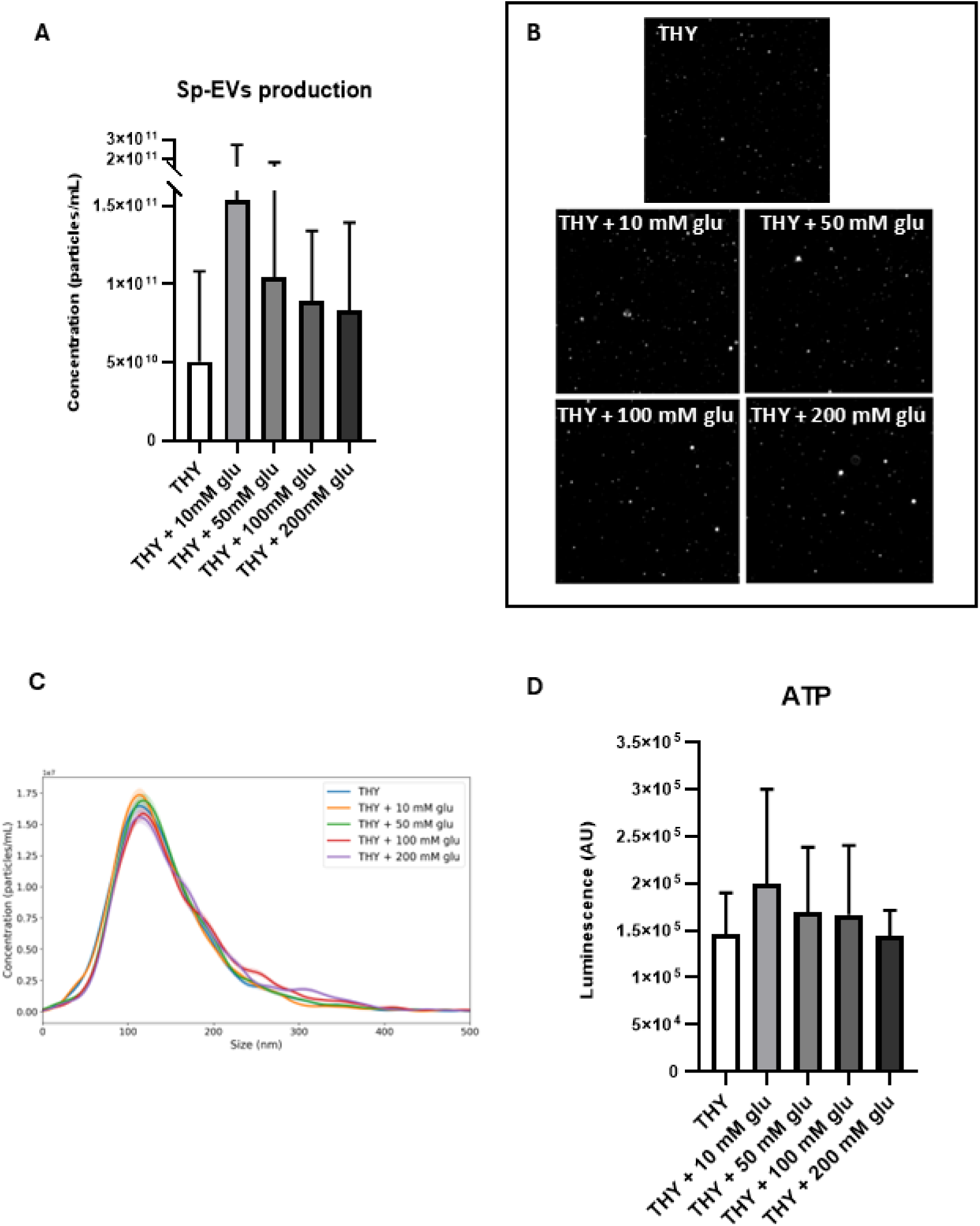
Glucose influence on Sp-EVs formation and ATP production by *S. pneumoniae* D39. (**A**) Graph representing Sp-EVs concentration (particles/mL) after nanoparticle tracking analysis. EVs were isolated from the supernatant of *S. pneumoniae* D39 grown until mid-logarithmic phase at 37°C in THY with D-(+)-glucose (10, 50, 100 and 200 mM). A condition with *S. pneumoniae* D39 grown in THY without D-(+)-glucose was used as control. The detected Sp-EVs concentration was normalised to the bacterial culture OD600 measured before EV isolation. Mean ± SD. (**B**) Representative snapshots of Sp-EVs, visualized by dynamic light-scattering microscopy. (**C**) Histogram showing particle profile for concentration (particles/mL) and size (nm) of Sp-EVs in all tested conditions. (**D**) Graph representing ATP production in *S. pneumoniae* D39 cells grown in THY with or without D-(+)-glucose (10, 50, 100 and 200 mM) at 37°C until mid-logarithmic growth phase. The amount of ATP present and extracted from bacterial cells was measured as luminescent signal and expressed in AU. The detected luminescence signal was normalised to the OD600 of the bacterial culture measured before the ATP extraction. Mean ± SD.

Glucose metabolism is directly linked to ATP production in bacterial cells ^50^ and ATP levels are regulated according to cellular energy demands for key cellular processes ^55^. Therefore, intracellular ATP level was measured after growing pneumococci in the presence of glucose (**Fig. 4D**). ATP amount was higher in the 10 mM glucose condition compared to both control and other tested glucose concentrations. The similar trend observed for Sp-EVs production and intracellular ATP level in response to environmental glucose availability suggests that a low glucose concentration (10 mM) is favorable for *S. pneumoniae* and may promote EV formation.

### Sp-EVs influence intra- and interspecies biofilm formation

So far, our results have shown that *S. pneumoniae* EVs production is influenced by environmental factors, including pH and glucose availability, suggesting that Sp-EVs may play a role in mechanisms of bacterial metabolic adaptation. Bacterial EVs are also known to contribute to biofilm formation, facilitating structural development, nutrient transport, and intercellular communication^56–58^.

To investigate the role of Sp-EVs in biofilm development and to further assess the potential impact of Sp-EVs produced under glucose abundance, biofilm formation of *S. pneumoniae* D39 in the presence of Sp-EVs was evaluated (**Fig. 5A**). Pneumococci were incubated with increasing concentrations of Sp-EVs (C1 or C2), isolated from mid-exponential bacterial cultures in THY with (10, 50, 200 mM) or without glucose.

**Fig. 5.**
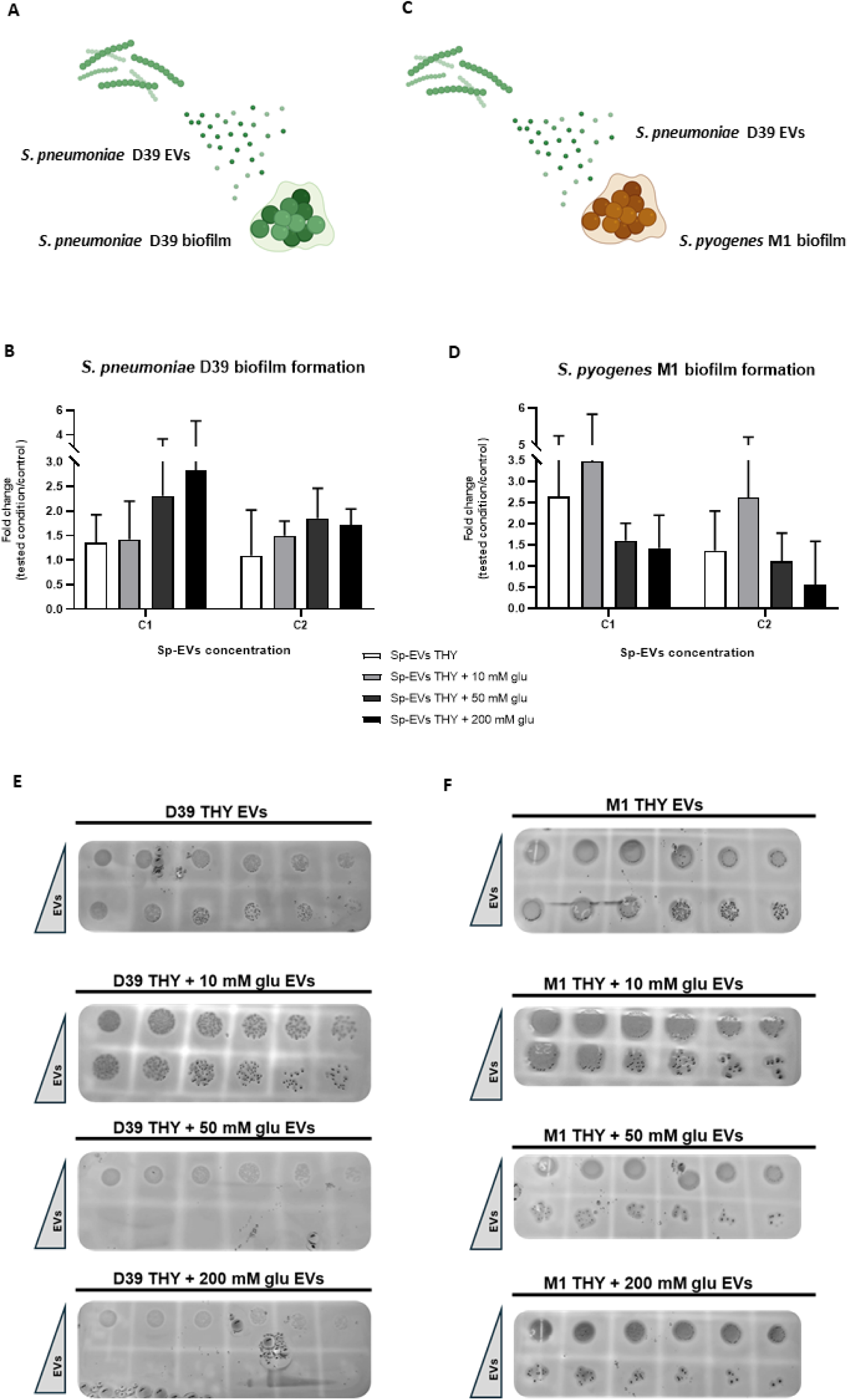
S**p-EVs influence on biofilm formation of *S. pneumoniae* D39 and *S. pyogenes* M1.** (**A, C**) Schematic representation of EVs effect on biofilm formation from different species. Quantification of biofilm formation by (**B**) *S. pneumoniae* D39 or (**D**) *S. pyogenes* M1 after incubation for 18 hours with Sp-EVs at two different concentrations (C1= ∼3 x 10^9^; C2= ∼4,8 x 10^10^). Sp-EVs were isolated from the supernatant of pneumococci grown until mid- exponential phase at 37°C in THY with or without D-(+)-glucose (10, 50 and 200 mM) and were subsequently diluted in THY. A condition of biofilm formation by bacterial cells without Sp-EVs was used as control. Fold change (tested condition/control) ± SD. (**E, F**) Representative images of spot assay performed to measure the viability of planktonic cells from the biofilm of *S. pneumoniae* D39 or *S. pyogenes* M1 after incubation with Sp-EVs for 18 hours as previously described. D39 or M1 bacterial cultures were serially diluted, spotted onto blood agar plates and grown at 37 °C overnight to evaluate CFUs.

Biofilm formation increased when pneumococci were grown for 18 hours in the presence of Sp-EVs, compared to the control (bacteria grown in the absence of Sp-EVs) (**Fig. 5B**). Moreover, pneumococcal biofilm development was particularly triggered by the lowest concentration of Sp-EVs (C1), isolated under the highest glucose concentrations (50 and 200 mM).

Next, we investigated whether Sp-EVs could also influence biofilm formation of a different streptococcal species, *Streptococcus pyogenes* M1 (**Fig. 5C**), another Gram-positive human pathogen that colonizes the throat, nose and skin^59,60^.

Similarly to the observed with the intra-species biofilms, Sp-EVs enhanced *S. pyogenes* M1 biofilm development, compared to the control without EVs (**Fig. 5D**). However, biofilm formation was tendentially higher when *S. pyogenes* M1 was exposed to Sp-EVs isolated from pneumococcal culture in THY without glucose or with a low glucose concentration (10mM). The viability of planktonic cells derived from the biofilm was also examined. Planktonic cell suspensions were serially diluted and spotted onto blood agar to determine the resulting colony forming units (CFUs). The viability of *S. pneumoniae* D39 planktonic cells decreased when pneumococci were incubated for 18 hours with the higher Sp-EVs concentration, as shown by a reduction in CFUs. The decrease in planktonic cells viability was even more pronounced when Sp-EVs at the higher concentration were produced under increasing glucose concentrations (10, 50, 200 mM) (**Fig. 5E**). A similar outcome was observed for *S. pyogenes* M1 biofilm, where the reduction of planktonic cell viability was directly correlated to the Sp-EVs concentration and glucose abundance, although a greater number of CFUs was detectable compared to the *S. pneumoniae* D39 conditions (**Fig. 5F**).

Overall, our results suggest that Sp-EVs promote biofilm formation in both *S. pneumoniae* D39 and a different streptococcal species, *S. pyogenes* M1. Interestingly, EVs produced under glucose abundance, have opposing effects on intra- vs inter-species biofilm formation.

## Discussion

The biogenesis and biological effects of extracellular vesicles (EVs) produced by the Gram-positive bacterium *Streptococcus pneumoniae* are still poorly understood. This study provides new insights into the environmental influence and functional roles of pneumococcal EVs (Sp-EVs), highlighting their potential role in bacterial metabolic adaptation and intra- or interspecies communication.

Bacteria constantly adapt to environmental conditions to ensure survival and proliferation ^1,2^. *S. pneumoniae* is a highly adapted human pathogen, capable of colonizing and invading its host, causing localized infections like otitis media and sinusitis, or more severe and life-threatening diseases like community-acquired pneumonia, bacteremia or meningitis^61^. The study of metabolic adaptation mechanisms in *S. pneumoniae* is essential for understanding its strategies for coping with the host environment. Bacterial regulatory systems are involved in sensing and responding to environmental signals, through the expression of virulence factors and proteins required to react to stressors such as antibiotics, temperature and pH variation and nutrient depletion^61^.

We demonstrate that human serum impacts the pneumococcal cell wall structure and modulates Sp-EV formation (**Fig. 1**). This suggests that host-derived factors, such as human complement proteins, may induce remodeling of the bacterial cell wall, potentially facilitating EV release. Factor H, a human protein involved in complement regulation, protects the septum of the pneumococcus against C3 binding and consequently opsonization^62^. The septum is a highly dynamic subcellular location, where the cell wall is constantly being remodeled and thus a vulnerable place for host attack. In line with this, proteomic analysis revealed an abundance of septum-related proteins in Sp-EVs (e.g. DivIVA, EzrA, FtsA, FtsE, FtsX, FtsZ), suggesting a possible septal origin for EV formation (**Fig. 3**). Given the fact that WGA staining was enriched in the septal region and higher EV production occurred upon NHS incubation, we hypothesize that the membrane attack complex (MAC), also termed C5b-9, at the terminal complement cascade, might be inserted into the cell wall of bacteria, leading to pore formation and thus facilitating the observed higher release of EVs. We optimized a protocol for NHS- pneumococcus incubation and observed C5b-9 deposition on the bacterial surface on a time- and concentration-dependent manner (**Fig. S1**). Moreover, *S. pneumoniae*-derived serum (after NHS incubation and removal of bacteria) showed marked reduction of C5b-9 deposition on CHO cells and less hemolysis of red blood cells, arguing for a depletion of C5b-9 from the serum, probably due to binding to bacteria (**Fig. S2**). These results point towards a possible role of MAC assembly and the release of EVs. More studies should be conducted to follow up.

During its progression from the nasopharynx to other niches of the human host, *S. pneumoniae* must also adapt to changes in temperature. It has been shown that during pneumococcal spread, the temperature can range from 30–32°C in the anterior nasopharynx, 37°C in deeper tissues as the lungs, bloodstream, and central nervous system and up to 40°C in feverish episodes^63^. Our data show that EVs production by *S. pneumoniae* D39 is sensitive to changes in temperature and it is enhanced at 37°C. On the other hand, our results highlight that a pH shift negatively affects Sp-EVs production: exposure to either basic or acidic environments slows down pneumococcal EV formation (**Fig. 2**). The effect of high temperatures on bacterial EVs formation was also observed in *Staphylococcus aureus*, where not only more EVs were produced but also the cargo was altered^64^. Interestingly, one of the proteins enriched in the Sp-EVs proteome is glutamate dehydrogenase (GdhA), a protein recently shown to play crucial roles in pneumococcal thermoadaptation as well as for nutrient metabolism and virulence^63^. The presence of GdhA in EVs suggests a possible redistribution of key metabolic enzymes under certain environmental conditions.

Proteomic analysis also revealed that Sp-EVs are particularly enriched in proteins associated with carbon metabolism. The presence of so many essential metabolic enzymes in EVs is a remarkable and unprecedent observation. Compelled by the presence of pneumolysin in the vesicles, in our previous study we investigated the hemolytic capacity of Sp-EVs but could not observe lysis at the used EVs-concentration^38^. The question remains if other enzymes in Sp-EVs have catalytic activity. Gram-negative bacteria-derived outer membrane vesicles (OMVs) have shown bactericidal activity due to the presence of one amidase in the vesicles^65^. EVs could be taken as *metabolic satellites*, protecting, distributing and delivering functional enzymes across cells and organisms.

Intriguingly, a lower glucose concentration (10 mM) promoted Sp-EVs production (**Fig. 4**). *S. pneumoniae* relies on glucose as the primary carbon source for survival and growth^66^. Glucose is the prevailing sugar in blood and inflamed tissues and it is known to regulate the expression of virulence factors such as cell wall associated proteins^67^. Therefore, Sp-EVs production might be upregulated in favorable conditions, where glucose is not present in excess^52^. Under glucose-limited conditions, such as those encountered during early colonization or in certain tissue niches, the increase in EV production might serve as signalling to neighbouring cells or even as local nutrient scavenging systems. More studies on the cargo of Sp-EVs under glucose growing conditions are needed.

Finally, we demonstrate that Sp-EVs influence both intra- and interspecies biofilm formation, suggesting that these vesicles play a role in intercellular communication (**Fig. 5**). It has already been reported that OMVs produced by *Helicobacter pylori*, a major etiologic factor in gastritis, act as a scaffold for cell-cell binding and increase biofilm formation on gastric epithelial cells^68^. In mixed *Streptococcus mutans - Candida albicans* biofilms, *S. mutans* produces EVs containing a factor that contributes to *C. albicans* biofilm formation^69^. We show that Sp-EVs enhance biofilm formation not only in *S. pneumoniae* but also in a different species, *S*.

*pyogenes*, which can colonize the upper respiratory tract. However, Sp-EVs isolated under glucose-rich conditions promoted *S. pneumoniae* biofilm formation, whereas Sp-EVs from glucose-poor conditions increased *S. pyogenes* biofilm. This divergent effect on the two species may be linked to the Sp-EVs cargo, which could differentially influence biofilm development. *Escherichia coli* EVs were found to inhibit group A *Streptococcus pyogenes* (GAS) peptidoglycan remodeling, inducing cell division defects and ultimately growth inhibition. Moreover, altered expression of many virulence genes has been reported in bacteria treated with *E. coli* EVs^70^. Although the mechanism remains unknown, Sp-EVs appear to serve as a platform for bacterial communication, likely delivering essential factors that promote biofilm development.

Taken together, our study supports that EV formation is not solely a byproduct of bacterial physiology, but a regulated process closely linked to the metabolic and environmental status of the cell. EVs are undoubtedly great delivery systems, but their active role on shaping bacterial metabolism deserves further attention. In the future, the intricate regulation of Sp-EVs formation should be dissected and their role as metabolic decoys explored.

## Material and Methods

### Bacterial strains and growth conditions

Two bacterial strains were used in this study: *Streptococcus pneumoniae* D39 and *Streptococcus pyogenes* M1^71,72^. Strains were grown in liquid Todd- Hewitt broth (Roth^®^) supplemented with yeast extract (THY) at 37°C with 5% (vol/vol) CO2. Blood agar plates were prepared from Blood agar (VWR^®^) with addition of 5% (vol/vol) defibrinated sheep blood (Thermo Scientific^®^). Growth was monitored by measuring the optical density at 600 nm (OD600).

### Super- Resolution Structured Illumination Microscopy (SR-SIM) for bacteria interaction with Normal Human Serum (NHS)

For interaction of *S. pneumoniae* with normal human serum (NHS), blood was taken from two healthy voluntary participants (who were not vaccinated against *S. pneumoniae*) in BD Vacutainer^®^ Clot Activator Tubes (BD Biosciences). After 15 minutes of incubation, the blood was centrifuged at 4,000 x g for 15 minutes at room temperature (RT). The serum (referred to as normal human serum, NHS) was separated from the clot, kept at -20 °C and defrosted only shortly before experiments.

To study interaction of *S. pneumoniae* with NHS, Super- Resolution Structured Illumination Microscopy (SR-SIM) was performed. *S. pneumoniae* D39 was grown in suspension (≈10^8^ CFUs) and pelleted at 7,200 x g for 3 minutes at 4 °C. The bacterial pellet was resuspended in 500 µL of serum and incubated for 30 minutes at 37 °C and 5% CO2. After incubation, pneumococci were separated from the serum by centrifugation at 7,200 x g for 3 minutes at 4 °C.

Bacteria were stained with a mixture of 4’,6-Diamidino-2-Phenylindole, dihydrochloride (DAPI) (1:500) and wheat germ agglutinin (WGA) CF^®^488 (1:1000) in phosphate buffered saline-Tween (PBST) for 1 hour at RT. Bacterial cells were then washed and fixed with 4% (vol/vol) paraformaldehyde (PFA) solution for 15 minutes at 4°C.

For the SR-SIM imaging, 10 µL of the sample was spotted on 1% (vol/vol) agarose pads. Agarose pads were covered with No. 1.5 coverslips (Roth^®^) and stored at 4 °C for further imaging. The SR-SIM data were acquired on an Elyra 7 system (Zeiss) equipped with a 63×/1.4 NA Plan-Apochromat oil-immersion DIC M27 objective lens (Zeiss), a Piezo stage, and a PCO edge sCMOS camera with 82% QE and a liquid cooling system with 16-bit dynamic range. Using Lattice SIM mode, images were acquired with 13 phases. WGA CF^®^ 488 was detected with a 488-nm laser and a BP 495–590 emission filter and DAPI was detected with a 405-nm laser and a 477/35 emission filter. Super resolution images were computationally reconstructed from the raw data sets using default settings on ZenBlack software (Zeiss) Images were analyzed using the Fiji ImageJ software^73^ .

### Extracellular vesicles isolation

For bacterial extracellular vesicles (EVs) isolation, *S. pneumoniae* D39 strain was grown on a solid blood agar plate overnight, and single colonies were inoculated into THY and grown at 37°C with 5% (vol/vol) CO2. To test various environmental conditions, *S. pneumoniae* was alternatively grown at 30° C, at different pH conditions (pH 5 and 8) and in presence of sugar or NHS. For this, THY was further supplemented with acetic acid (0.6 g/mol) (Sigma-Aldrich^®^) or sodium hydroxide (0.4 g/mol) (Sigma-Aldrich^®^), D-(+)-glucose (10, 50, 100 and 200 mM) (Merck^®^) or NHS. At mid-logarithmic growth phase, 50 mL aliquots of bacterial culture were centrifuged twice (10,000 x g, 20 minutes, 4°C and 4,000 x g, 20 minutes, 4° C). The pellet was discarded, and the supernatant was filtered through a 0.45-μm pore membrane (Sartorius^®^) in order to obtain cell-free supernatant. The resulting cell-free media was ultracentrifuged at 100,000 x g for 2 hours at 4°C (Rotor SW 45 Ti, Optima XPN-80 Ultracentrifuge, Beckman Coulter, Life Sciences). The resulting vesicle pellet was resuspended in 1 mL sterile Milli-Q water and stored at 4°C for further analysis. For the environmental influence analysis, EVs were alternatively precipitated using ExoQuick-TC (System Biosciences) according to the manufacturer’s protocol. For subsequent use, EVs were stored at -80°C and thawed when needed.

### Extracellular vesicles counting

To determine size and concentration of isolated vesicles, nanoparticle tracking analysis was performed using the ZetaView^®^ (Particle Metrix GmbH) with a detection wavelength at 488- nm in scatter mode equipped with ZetaView software (version 8.05.16 SP3). Isolated vesicles were dispersed in 1 mL of Milli-Q water and diluted suitably.

For the experiments testing the environmental influence, isolated vesicles were counted using an NS300 dynamic light-scattering microscope (Malvern Panalytical) fitted with NanoSight NTA 3.2 software. Videos were captured at 24 fps for three periods of 60 seconds for each sample and analyzed using NanoSight NTA 3.2.

### Protein-protein interaction network analysis

A protein-protein interaction (PPI) network was constructed based on previously acquired proteomic data^38^ to identify key functional relationships among differentially expressed proteins. First, proteins identified from *S. pneumoniae* D39 strain were mapped to the well-annotated R6 reference strain identifiers using BLASTp^74^ with default parameters. A summary of statistics of the mapping is provided in Supplementary Table 1 (**Table S1**). Subsequently, the mapped R6 protein identifiers were used to construct the protein interaction network using the STRING database^75^. Protein interactions were inferred based on integrated evidence from multiple sources, including experimentally validated interactions, curated databases, co-expression data, genomic context predictions, and text mining, with a confidence threshold of 0.7. Network visualization was performed using Cytoscape^76^. Functional enrichment analysis of the network was conducted using KEGG pathway annotations to reveal biological processes and pathways associated with the identified protein interactions. Over-representation was evaluated using hypergeometric tests and Benjamini–Hochberg correction for multiple comparisons. Pathways with an adjusted p-value ≤ 0.05 were considered significantly enriched.

### ATP measurement

To quantify ATP in bacterial cells grown in the presence of sugar abundance, *S. pneumoniae* D39 was grown in THY with or without D-(+)-glucose (10, 50, 100 and 200 mM) (Roth^®^) at 37°C with 5% (vol/vol) CO^2^ until mid-logarithmic growth phase was reached. Bacteria were seeded in a 96-well plate and the BacTiter-Glo^™^ Microbial Cell Viability Assay was performed according to the manufacturer’s protocol. In this assay, luminescence is the result of mono- oxygenation of luciferin catalyzed by luciferase in the presence of Mg2^+^, ATP and molecular oxygen. The measured luminescent signal is proportional to the amount of ATP present and extracted from bacterial cells. The detected luminescent signal was normalized to the OD600 of the bacterial culture measured before the ATP extraction.

### Biofilm quantification

For quantification of biofilm formation, the Microtiter Dish Biofilm Formation Assay was performed with small changes^77^. Bacteria (OD600 0,3) were incubated with pneumococcal EVs at two different concentrations (C1 and C2) and statically grown on 96-well plates to obtain biofilms (Thermo Scientific), for 18 hours at 37°C with 5% (vol/vol) CO2. Four different EVs conditions were tested: EVs isolated from *S. pneumoniae* D39 culture in THY with or without D-(+)-glucose (10, 50, and 200 mM). After 18 hours, the supernatant was transferred to another plate and OD600 measured as a read for planktonic growth. To each well of the original plate, 200 µL of a 1% (vol/vol) crystal violet solution was added and the plate was incubated for 30 minutes at room temperature. After repeated washing steps, the plate was left to dry for 1 hour.

Ethanol [95% (vol/vol)] was added to each well and left for 30 minutes at room temperature. The released crystal violet was finally transferred to a new 96-well plate, and absorbance at 620 nm was measured.

### Spot assay

Bacterial cells were incubated in a 96-well plate with pneumococcal EVs at 37°C with 5% (vol/vol) CO2 for 18 hours to test biofilm formation as previously described. After 18 hours, planktonic cells suspension was transferred to another plate and OD600 was measured. For spot assay, planktonic cells were serially diluted, spotted onto blood agar plates and grown at 37 °C and 5% (vol/vol) CO2 overnight. Four different EVs conditions were tested: EVs isolated from *S. pneumoniae* D39 culture in THY with or without D-(+)-glucose (10, 50, and 200 mM).

### Statistical analysis

Unless otherwise stated, statistical significance was determined by ordinary one-way analysis of variance (ANOVA) test with a Tukey multiple comparisons test. Probability values (p- values) were defined as follows: ns, p ≥ 0.1; *, p ≤ 0.05; **, p ≤ 0.01; ***, p ≤ 0.001; ****, p ≤ 0.0001. Statistical analysis was performed using Prism version 9 for Windows (GraphPad Software, La Jolla, CA, USA).

## Acknowledgements

We thank the Microverse Imaging Center (and Aurélie Jost / Patrick Then) for providing microscope facility support for data acquisition (and data analysis). The ELYRA 7 (used for producing images on Fig.1) was funded by the Free State of Thuringia with grant number 2019 FGI 0003. The Microverse Imaging Center is funded by the Deutsche Forschungsgemeinschaft (DFG, German Research Foundation) under Germanýs Excellence Strategy – EXC 2051 – Project-ID 390713860. This work was further supported by the Bundesministerium für Bildung und Forschung (BMBF) within the program Computational Life Sciences, project MuMoSim (FKZ: 031L0291A).

## Author Contributions

MB and CV conceived and designed the experiments. MB, TF, LZ, LT and CV performed the experiments. MB, YB, CS, TF, LZ, LT and CV analyzed the data. CV and MTF contributed with reagents, materials, and/or analysis tools. MB, LZ and CV wrote the manuscript. MB, CV, YB, CS, MTF corrected the manuscript.

